# Environmental Enrichment Normalizes Hippocampal Timing Coding in a Malformed Hippocampus

**DOI:** 10.1101/130450

**Authors:** Amanda E. Hernan, J. Matthew Mahoney, Willie Curry, Greg Richard, Marcella M. Lucas, Andrew Massey, Gregory L. Holmes, Rod C. Scott

**Affiliations:** Department of Neurological Sciences, University of Vermont College of Medicine, Burlington, Vermont, USA 05405; Department of Neurology, Geisel School of Medicine at Dartmouth, Lebanon, New Hampshire, USA 03756; University College London, Institute of Child Health, London WC1N 1EH, UK

**Author notes:** Corresponding authors: Rod Scott, 95 Carrigan Drive, Stafford 118D, Burlington, VT 05405, Amanda Hernan, 95 Carrigan Drive, Stafford 118F, Burlington, VT 05405. The authors contributed equally to this work (Amanda E. Hernan, J. Matthew Mahoney). MCD: malformation of cortical development, EE: environmental enrichment, FST: fine spike timing, ACG: autocorrelogram.

**Keywords:** cortical malformations, environmental enrichment, cognition, temporal coding, rate coding, hippocampus

## Abstract

Neurodevelopmental insults such as malformations of cortical development (MCD) are a common cause of psychiatric disorders, learning impairments and epilepsy. Animals with MCDs have impairments in spatial cognition that, remarkably, are improved by post-weaning environmental enrichment (EE). To establish the network-level mechanisms responsible for these impacts, hippocampal *in vivo* single unit recordings were performed in freely moving animals in an open arena. We took a generalized linear modeling approach to extract fine spike timing (FST) characteristics and related these to place cell fidelity used as a surrogate of spatial cognition. We find that MCDs disrupt FST and place-modulated rate coding in hippocampal CA1 and that EE restores both to normal. Moreover, FST parameters *predict* spatial coherence of neurons, suggesting that mechanisms determining FST are critical for cognition. This suggests that FST parameters could represent a therapeutic target to improve cognition even in the context of a structurally abnormal brain.

**HIGHLIGHTS:**

- Environmental enrichment (EE) in rats with cortical malformations improves cognition.
- EE resolves impaired rate and timing coding of hippocampal pyramidal neurons.
- Taken together, circuit-level dynamics directly affect quality of the cognitive map.

**RESEARCH IN CONTEXT:** Insults during neurodevelopment, particularly those that result in physical malformations in the brain, lead to cognitive impairment, psychiatric disorders and epilepsy. Environmental enrichment (EE) improves cognitive outcome in patients and animal models with brain malformations. Understanding how EE can improve cognition at the level of neural networks can lead to new treatment targets. Remarkably, using an approach that mathematically models neuron firing we show that firing is mistimed in animals with malformations and that EE improves this abnormality. Importantly, timing abnormalities predict abnormalities in cognition at the single neuron level, suggesting that restoring timing could improve learning and memory deficits.

## INTRODUCTION

Neurodevelopmental insults are common etiologies in many neurological diseases, including epilepsy, schizophrenia, cerebral palsy, and autism spectrum disorders. Malformations of cortical development (MCDs) are one result of an insult during neurodevelopment and are commonly identified in patients with significant cognitive impairments and seizures (Kuzniecky et al. 1993). Cognitive impairments are a major driver of impaired quality of life and therefore strategies that maximize cognition would be expected to have a major positive impact on outcomes. In the context of epilepsy, the main therapeutic strategies have targeted seizures in order to try to achieve cognitive improvements. Unfortunately, this approach has had limited success, raising the issue of whether abnormal neural networks underlying cognitive impairments can be functionally modified to improve cognition independently of treating seizures. We have previously addressed this issue behaviorally in the methylazoxymethanol (MAM) model of MCDs in rats. *In utero* injection of methylazoxymethanol acetate (MAM) produces offspring with neuropathology similar to patients with MCDs associated with early onset epilepsy. These animals display significant impairments in spatial learning and memory across two different behavioral tasks(Jenks et al. 2013; Lucas et al. 2011). Remarkably, these impairments are dramatically improved following environmental enrichment(Jenks et al. 2013), suggesting that brains with MCDs can be functionally altered to improve behavior. However, the nature of the underlying neural network changes responsible for this improvement is unknown. Extrapolating from environmental enrichment to a broader class of therapies targeting cognitive impairment in MCDs is impossible, given that there is no obvious target. Characterization of the system-level mechanisms of improvement following environmental enrichment could provide critical insight for development of novel therapies for improving cognition in patients with MCDs.

Information processing in the brain is a function of neural dynamics, conceptualized here as system-level mechanisms. The hippocampus is a multi-layered series of integrated networks that underlie spatial cognition(Eichenbaum et al. 1999) and therefore modifications to the hippocampal neural system in MAM exposed animals with MCDs could explain both the spatial cognition impairments and the observed improvements following environmental enrichment. Pyramidal cell action potential firing within these networks is precisely modulated both in space and in time with respect to neuronal oscillations in the local field potential(Muller 1996). Modulation of firing rate (rate coding) with respect to spatial location is observed with hippocampal place cells. Modulation of action potential firing with respect to time (timing coding) is observed in the phenomena of phase preference and phase precession(Skaggs et al. 1996). The rate and timing modulations of pyramidal cell firing dynamics are sculpted by a wide variety of interneurons, including those targeting the axon initial segment and predominantly expressing parvalbumin. This precise modulation of pyramidal cell firing with respect to space and time is crucial for information processing, resulting in effective spatial learning and memory(Dragoi & Buzsáki 2006). We therefore hypothesize that MCDs are associated with abnormalities in rate and timing coding in hippocampal CA1 pyramidal cells, and that environmental enrichment, which is therapeutically effective in this model, will recover these functional abnormalities, even in the context of a malformed brain.

Both rate and timing coding influence the probability of an action potential (*spike*) at any given time. Typical summaries of spike trains, such as autocorrelograms (ACGs), are complex mixtures of all effects that dynamically modulate firing. Therefore, to dissociate rate and timing effects from each other, we used a generalized linear modeling approach that incorporates both a neuron’s spatial receptive field and its timing properties. The GLM approach is a statistically principled way to extract structure from spike trains that explicitly models timing coding over-and-above spatial rate coding. The GLM captures timing features robustly without, for example, requiring a minimum firing rate for analysis. Using the GLM, we are able to extract timing features that can then be quantitatively related back to rate coding parameters without *ad hoc* summaries of spiking dynamics or conceptual circularity inherent in trying to examine temporal coding from the ACG alone.

We show for the first time that rate and timing coding in hippocampal CA1 pyramidal cells is disrupted in animals with MCDs and that this disruption is associated with a reduction in parvalbumin expressing interneurons. Environmental enrichment modifies rate and timing modulation disruptions and increases the number of parvalbumin expressing interneurons, suggesting that structurally abnormal networks can be functionally restored towards normal without alteration of the gross structural abnormality. This supports the view that strategies that normalize pyramidal cell firing dynamics have substantial therapeutic promise for patients with MCDs.

## MATERIALS AND METHODS

### Animals

All animal procedures were approved by the Dartmouth College Institutional Animal Care and Use Committee and University of Vermont Institutional Animal Care and Use Committee, under United States Department of Agricultural and Association for the Assessment and Accreditation of Laboratory Animal Care International approved conditions, in accordance with National Institutes of Health guidelines. Sprague-Dawley rats housed with a 12 h light/dark cycle and *ad libitum* access to food and water.

### MAM model

Dams were randomly selected for intraperitoneal injection with either 20 mg/kg MAM (Midwest Research Institute Global, Kansas City, MO) or saline at embryonic day (E) 17.

### Enrichment paradigm

Pups were raised with their dams until weaning at postnatal day (P) 25, when they were then randomized to either standard or enriched housing until electrophysiological studies were performed. Males and females were included. Animals that were enriched were housed in breeder cages (38 x 48 cm) with 1 to 3 other animals as well as various colorful and differently shaped and textured inanimate objects. Animals that underwent standard care were housed singly in a rat cage with no inanimate objects. All animals stayed in their assigned environment throughout the behavioral experiments and underwent only the reported treatments, behavioral and electrophysiological experiments until they were sacrificed for histology. Due to the nature of the experimental setup, experimenters could not be blinded to group. 15 rats were used: 4 controls (59 cells), 7 MAM unenriched (101 cells), and 4 MAM-E (98 cells).

### Surgical implantation of electrodes

Rats were anesthetized with isoflurane (2-3 % in oxygen) and placed in a stereotaxic frame (Kopf Instruments, Tujunga, CA). Custom-built electrodes containing four independently drivable tetrodes were implanted 3.2 mm posterior to the bregma, 2.2 mm lateral and 1.7 mm deep into the dorsal CA1 hippocampus.

### Data Acquisition

Tetrode assemblies were advanced 50 μm twice a day until hippocampal theta oscillations (6-12 Hz), sharp waves and ripples were observed in the EEG. Electrodes were then advanced in 25 μm increments until CA1 single unit activity was detectable. Single unit activity was recorded when waveforms above 40 μV in amplitude were observed on one or more tetrodes. The signal from the electrodes was preamplified directly from the rat’s head by operational amplifiers and transmitted via a custom cable to a Neuralynx recording system (Neuralynx, Bozeman, MT). Recordings were performed in an open arena. The arena had a diameter of 75 cm and the walls and floor were made of plywood and painted grey. To provide a visual anchoring cue, a white piece of paper covering 100° of arc was attached to the wall of the arena. Rats were allowed to freely explore the arena for 15 min per session per day. To avoid recording the same cells twice, the tetrodes were advanced at the end of each session. The animals were tracked in concurrent video recordings so that cell firing relative to position could be analyzed.

### Electrophysiological analysis

Action potentials were clustered using Neuralynx Spike Sort 3D software (Bozeman, MT). Once cells had been clustered, firing rate maps were computed in MatLab (MathWorks, Natick, MA) showing action potential firing as a function of position. We calculated place fields by smoothing spatially binned spike counts and occupancy maps (time spent in each pixel) using a 2D Gaussian kernel. We chose the width of the kernel using 5-fold cross-validation with the log-likelihood of the resulting place field model as the measure of model fit. Due to the recording protocols, very few interneurons were recorded over all. Those that were recorded were identified as distinct from pyramidal cell groups in that they typically do not exhibit place preference and have a much higher rate of firing than pyramidal cells. Interneurons or neurons that did not appear to be hippocampal in origin were excluded from further analysis.

Spatial coherence of pyramidal cells was calculated for the entire population as previously described. Coherence is defined as the correlation (2D z-correlation) between the rate in each pixel and the average rate of its 8 nearest neighbors (Lenck-Santini & Holmes 2008; Liu et al. 1994). The peak rate of a field corresponded to the average rate in the highest firing pixel and its neighbors. Most analyses were conducted on the entire pyramidal cell population; for comparison purposes, “classical place cells” were defined as pyramidal cells with a spatial coherence of 0.3 or greater, as previously reported (Lenck-Santini & Holmes 2008). Inclusion of a given recording session required adequate sampling of the environment and place cell coherence was evaluated based on speed-filtered sessions. Mean firing rate was calculated as the number of action potentials divided by the total time of the recording session. Instantaneous firing rate was calculated as 1/peak interspike interval. Spike width was calculated by locating the first point and the peak of the waveform, and then finding a point that was 25% maximal height and calculating the time between the two measures.

### Histology

102 sections from 8 rats: 2 control (4-17 sections), 4 MAM (5-15 sections), 2 MAM-E (8-13 sections) were used. Animals were deeply anesthetized with isoflurane (4-5% in oxygen) immediately after which rats were perfused transcardially with 1x PBS followed by 4% paraformaldehyde (Sigma Aldrich, St. Louis, MO). Brains were postfixed in 4% paraformaldehyde overnight, 30% sucrose embedded until they sank and frozen in OCT tissue embedding solution. Coronal sections (40 μm) through the entire extent of the hippocampus were collected using a cryostat (Leica CM3050, Leica Microsystems). Before immunostaining, sections were washed 2 times in PBS for 15 minutes. They were then placed in a blocking buffer solution composed of 1x PBS, 0.5% Triton, and 10% normal goat serum for 1 hour at room temperature. After blocking, sections were incubated in mouse anti-parvalbumin primary antibody (1:500, Millipore cat # MAB1572, RRID: AB_2174013) for 18 hours at 4°C. Sections were once again washed twice in PBS for 15 min per wash. They were then incubated in species-specific secondary antibody (AlexaFluor 488, Life Technologies) overnight, again at 4°C. After two 10 minute washes in PBS, sections were lastly incubated in DAPI solution for an additional 10 minutes. Sections were mounted on charged slides (Superfrost Plus, Fischerbrand) and coverslipped with mounting medium (Fluoromount, Sigma).

### Generalized linear models

We binned pyramidal cell spike trains into ◻ = 1 ms bins and fit generalized linear models (GLMs) to the resulting binary sequence. The GLM had two time-varying components: the place field of the pyramidal cell, *p*(*x*, *y*), where (x,y) is the 2D position of the rat, and the post-spike filter (PSF), *f* (*s*), which models the post-spike rate modulation as a function of the time, s, after a spike.

For the PSFs, we used ten ‘raised cosine bump’ basis functions, *b*_*i*_(*s*) (i = 1,…,10), extending over 0.7 s and one 1 ms impulse, *b*_11_(*s*), immediately post-spike to capture the absolute refractory period after a spike (cf.(Pillow et al. 2008)). Thus, the PSF is defined by a set of 11 coefficients, *◻*, for these 11 basis functions:

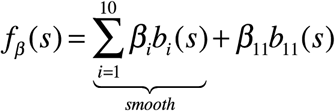

where we call the first term on the left hand side corresponding to the raised cosine basis the *smooth component* of the PSF. The conditional intensity function, *◻*(*t*), of the full model at a given choice of *◻* is given by

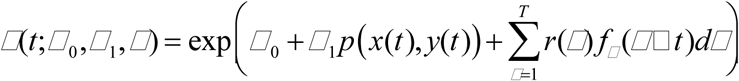

where *r*(*t*) denotes the spike train, *◻*_0_ controls the baseline firing rate of the neuron, *◻*_1_ modifies the firing intensity of the place field, *f*_*◻*_ is the PSF defined by *◻*, and T denotes the length of the recording.

We fit the GLM by maximum likelihood with a ridge penalty. The penalized likelihood function is

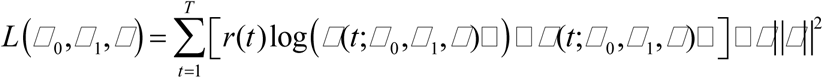

where *◻* is the penalty hyperparameter, which corresponds to the variance of a Gaussian prior on the PSF coefficients. We selected *◻* from a grid by evidence maximization as in (Park et al. 2014). Note that we only penalize *◻* the coefficients, treating the place field and background firing as covariates.

### Clustering post-spike filters

We normalized the PSFs to have a total power of one and clustered the normalized PSFs using k-means clustering (k = 2). We chose k = 2 by visual inspection of the clustered correlation matrices, and the heat map of clustered PSFs (Figure 2). The larger of the two clusters contains cells that are strongly up-regulated post-spike, which we call *bursty cells*.

### Analysis of post-spike filter parameters

We analyzed six quantitative features of the PSFs that captured distinct aspects of temporal coding. We first reduced each PSF to its smooth component (see above). For a PSF *f*, we defined *rate normalized power* as 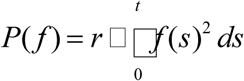, where *r* is the mean firing rate of the cell and *t* is the duration of the PSF (∼0.7 s). This normalization was chosen because the integral of the squared PSF (*raw power*) is strongly negatively correlated to the firing rate (data not shown). This is likely due to the fact there is a relatively constant amount of total firing rate modulation per cell, so low firing rate cells attribute more modulation *per spike* and thus have higher raw power.

To study ‘shape’ properties that are independent of power, we normalized each PSF to have total power equal to one:

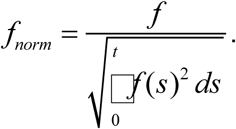

We defined burstiness from the immediate post-spike interval value as 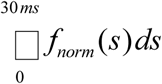. This value is highly correlated with the instantaneous frequency (defined as 1/peak interspike interval) as well as >30ms ISIs. For bursty cells (Fig. 1), we defined the *theta peak* as the maximum value of *f*_*norm*_ in the interval 0.083-0.167s (i.e. one theta cycle) post-spike. Likewise, the *theta trough* was the minimum value attained in the interval 0.042-0.083s, and *theta depth* was defined as theta peak minus theta trough. We defined *theta correlation* as the maximum linear correlation between *f*_*norm*_ and a theta-frequency cosine wave *g*_*A◻*_, (*t*) = *A*cos(*◻t*):

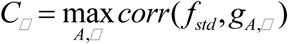

where the amplitude, *A*, is constrained to be positive and the frequency, *◻*, is constrained to be in the interval 6-12Hz.

**Figure 1.**
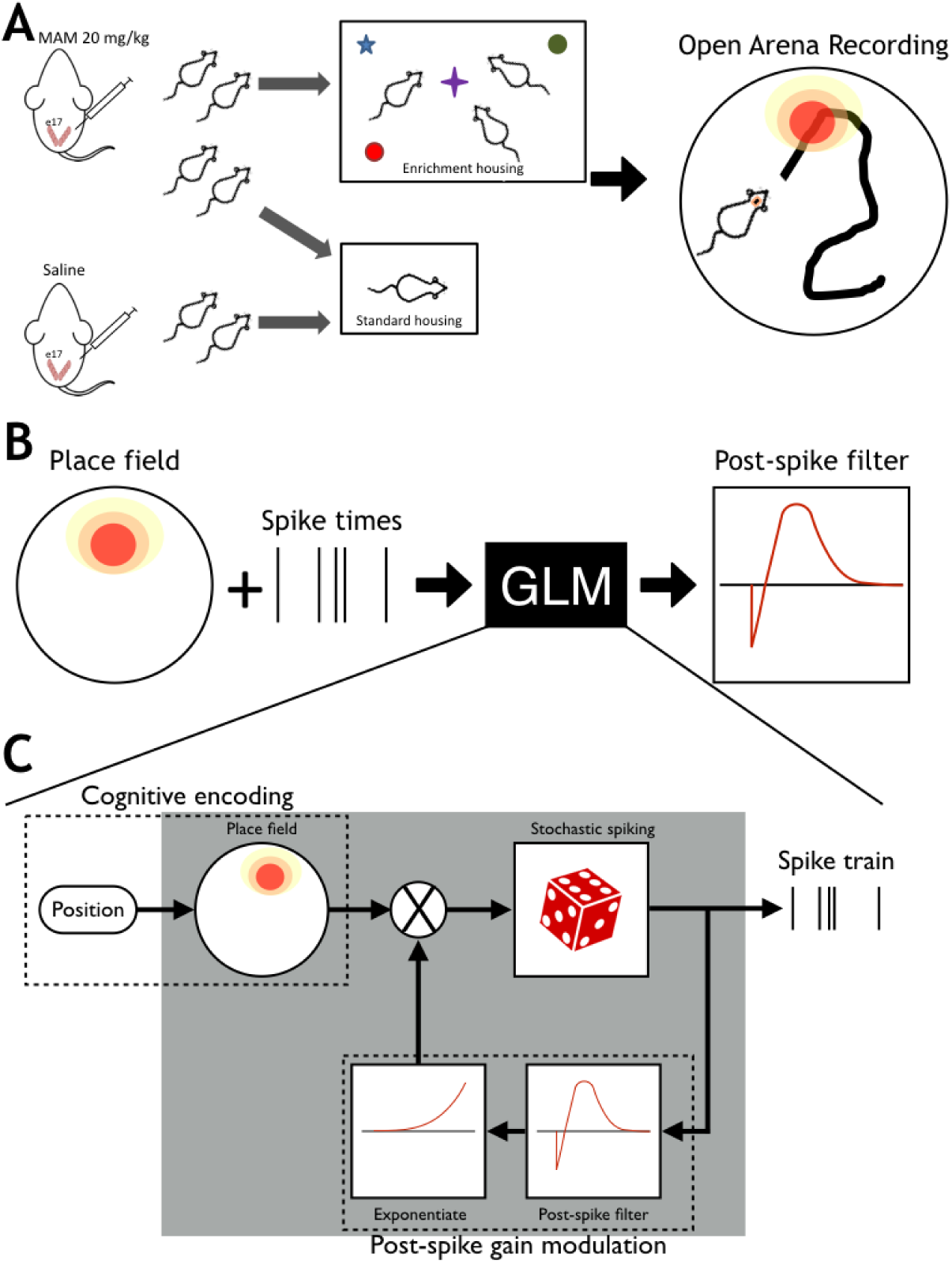
Environmental enrichment and analysis pipeline. (**a**) Pregnant dams were injected I.P. with methylazoxymethanol acetate (MAM) or saline (Control) at embryonic day 17. Post-weaning, control pups and one group of MAM pups were housed in standard conditions, while another group of MAM animals received environmental enriched (EE), including multiple novel objects to explore and other animals with which to interact. All animals were implanted as adults (>p60) with four tetrodes in CA1 and were recorded foraging in a familiar environment for food reward. Place fields were constructed by comparing spike times to the animal’s position acquired from video tracking. (**b**) Place fields of CA1 pyramidal cells and spike times were used in a generalized linear model (GLM) to model the fine-spike timing of cells, which is captured with a post-spike filter (PSF). Specifically, the PSF models the short timescale (< 650 ms) post-spike modulation of firing probability and captures the autocorrelation structure of cell firing. (**c**) As a stochastic model, the GLM considers the firing probability to be a function that varies in time. The cognitive component of the GLM uses the place field of the cell, which is a slowly varying, position dependent firing rate (top left). To capture the fast timescale structure, the GLM uses a PSF, which feeds back through an exponential nonlinearity to modulate the firing probability (bottom right). The PSF is defined by 11 parameters that are chosen to maximize the likelihood of observing the measured spike train. Thus, the PSF is learned through the GLM with input from the place field and the raw spike train. Note that the modulation by the PSF is multiplicative. Hence, the PSF corresponds to ‘gain modulation’. The PSF is parametrized by a set of basis functions. The fitted PSF is learned from the measured spike train (top right) through penalized maximum likelihood estimation.

### Statistical Analysis

We use generalized estimating equations (GEE) in SPSS (21.0 Chicago, III), which allows for within-animal correlations and the assumption of the most appropriate distribution for the data; all data distributions were visually assessed and the most appropriate link function was used. Goodness of fit was determined using the corrected quasi likelihood under independence model criterion and by the visual assessment of residuals. One degree of freedom was assumed based on a Wald chi-square within the GEE framework. We compared spatial coherence and PSF parameters across groups using GEE as multiple cells were obtained from individual animals. These cells are likely to be related to each other given that they are recorded in the same network. PSF shapes were compared between groups by down sampling the PSF to 10 ms bins across the entire timecourse of the PSF. Group by time interactions were compared to establish whether the PSFs all come from the same distribution. In addition to using *a priori* parameters of the PSF we also used a dimension reducing factor analysis in SPSS (21.0 Chicago, Ill) to identify factors in an unsupervised way. We obtained a Varimax rotated solution and generated factors for each cell for each solution. Factors with an eigenvalue of >1 were retained for further analysis. Cell counts were compared between groups using a negative binomial distribution appropriate for counts data. All values reported are estimated marginal means +/-SEM. Multiple regression analyses within GEE with spatial coherence as the dependent variable and PSF parameters (or in a separate analysis, factors) and 2-way interaction terms as independent variables. Predicted values from the analyses were retained. Spearman and Pearson correlations were used to determine the relationships between observed and expected data and to obtain an objective measure of how much variance was explained by the regression analyses.

## RESULTS

In order to understand the neural networks underpinning poor spatial cognition and improved cognition after environmental enrichment (EE) in animals with MCDs, we used *in vivo* electrophysiological tetrode recording of pyramidal cells in CA1 of the hippocampus (Figure 1). We monitored three groups of animals during foraging in an open field: 1) Controls, 2) MAM animals (MAM), and 3) MAM animals exposed to post-weaning environmental enrichment (MAM-E). To determine the effects of MCDs and EE on neural circuits, we studied 258 hippocampal CA1 pyramidal cells from 15 rats: 4 controls (59 cells), 7 MAM unenriched (101 cells), and 4 MAM-E (98 cells). Table 1 provides information regarding standard electrophysiological properties of all pyramidal neurons recorded in our cohort. As a whole population, the action potential half width (p=0.313), mean firing rate during the session (p=0.758) and instantaneous frequency (p=0.149) of firing were not different between the groups, indicating homologous populations of neurons were recorded from all groups of animals (Table 1).

**Table 1.** Description of the standard firing characteristics of all recorded populations of pyramidal neurons.

|  |  | Spatial Coherence | std. error | Spike Width | std. error | Mean Firing Rate | std. error | Instantaneous Firing Frequency | std. error |
| --- | --- | --- | --- | --- | --- | --- | --- | --- | --- |
| Bursty | Control | 0.30 | 0.023 | 0.32 | 0.017 | 0.48 | 0.109 | 25.27 | 4.693 |
|  | MAM | 0.22 | 0.012 | 0.35 | 0.022 | 0.77 | 0.163 | 25.45 | 2.568 |
|  | MAM-E | 0.32 | 0.034 | 0.36 | 0.030 | 0.76 | 0.026 | 36.48 | 3.751 |
| Refractory | Control | 0.12 | 0.034 | 0.30 | 0.023 | 1.08 | 0.304 | 10.52 | 1.744 |
|  | MAM | 0.10 | 0.023 | 0.31 | 0.026 | 0.52 | 0.206 | 5.30 | 0.726 |
|  | MAM-E | 0.21 | 0.037 | 0.36 | 0.047 | 1.55 | 0.444 | 9.97 | 1.625 |

### Spatial coherence of CA1 pyramidal cells is impaired in MAM and restored in MAM-E

We hypothesize that, consistent with previously observed behavioral impairments, animals with MCDs have impaired spatial rate coding in CA1 pyramidal neurons and that EE would ameliorate these abnormalities. This would suggest that EE modifies local networks to improve spatial cognition. We used *spatial coherence* (Muller 1996) to measure the quality of cognitive encoding by CA1 pyramidal neurons during open field foraging. Spatial coherence of pyramidal cell firing was significantly altered by MCD and recovered in the enrichment paradigm. Across all recorded pyramidal cells, coherence was 0.192 ± 0.008 in control animals and 0.145 ± 0.015 in MAM animals (p=0.005), consistent with our behavioral findings that MAM animals perform poorly in spatial tasks (Jenks et al. 2013). Environmental enrichment significantly increased spatial coherence of pyramidal cells to 0.256 ± 0.029 (p=0.001; MAM-E compared to MAM unenriched), consistent with our prior findings that EE significantly improves spatial learning and memory. These changes in spatial coherence can be seen best in Figure 7b, which shows the spatial coherence values as a population average (bar graphs, inset) and as a moving average of four by group. While the initial peak in the MAM and control groups are similar, the MAM group does not have the right “tail” seen in the control group, representing neurons with high coherence values. The MAM-E group has these two peaks, although the lower coherence peak is shifted to the right thereby increasing the average coherence of the whole population. Observations noted in the moving average plot can also be seen as a significant increase in the proportion of cells that met criteria for classical place cells (defined as pyramidal cell having a coherence of 0.3 or greater (Muller & Kubie 1987)) from 21% in MAM animals to 42% in MAM-E animals. These results demonstrate that EE restores the quality of spatial encoding despite the cortical malformation.

### Timing coding is heterogeneous across groups

The non-random firing of spikes within theta oscillations of the local field potential (LFP) and relative to each other (autocorrelation) is strongly associated with spatial cognition(Buzsáki 2002). These phenomena fall under the broad rubric of *timing coding*, and require microcircuits that can dynamically modulate firing probabilities of neurons. Because MAM has significant effects on neural anatomy, we hypothesize that MCDs result in damaged hippocampal circuits whose dynamics are suboptimal for encoding spatial information and that EE would restore these dynamics. To quantify the timing coding of neurons, we modeled spike trains using a generalized linear model (GLM). Mathematically, our GLM is a statistical model of the firing probability of a neuron that simultaneously incorporates the many tuning properties of spiking, including the spatial rate coding (the *place field*) and fine spike timing (FST) properties of the cell. The FST properties include all effects where the relative timing of spikes is different from predicted based on spatial rate coding alone This means that the autocorrelation of spiking is different from expectation when only a place field is used to model the spike train. FST results from the fact that the firing probability of a cell changes as a function of the history of the spike train i.e. the timing of a previous spike influences the current firing probability (Figure 1). The GLM captures FST properties using post-spike filters (PSFs), which are parameterized curves that describe the modulation of firing probability after a spike (Figure 1). The PSF, therefore, is a functional *endophenotype* encapsulating the intricate circuit dynamics controlling the high-speed modulation of the probability of spiking. We find that by adding the PSF to the GLM we can accurately account for the autocorrelation structure of neural firing.

To analyze the timing coding properties of each cell, we quantified the variation in the PSFs across the whole population. Our initial analyses sought to examine specific aspects of temporal coding known to be important for spatial cognition in CA1 pyramidal neuron firing. To make comparisons among cells, it was necessary to first normalize all PSFs to have total power equal to one (see Materials and Methods) since the *total power* of the PSFs, i.e. the integral of the squared PSF, varied substantially and was strongly correlated to firing rate (see discussion in Materials and Methods). We clustered the normalized PSFs using the k-means clustering algorithm (Onoda et al. 2007; Lloyd, 1982) and identified two distinct clusters of cells (Figure 2a). The first, and smaller, cluster contained cells (*refractory cells)* whose firing was slightly down-regulated immediately following a spike (negative value of PSF immediately after t = 0, Table 2). The second, and larger, cluster contained cells that were markedly up-regulated post-spike (*bursty cells*). The bursty cells also showed consistent theta modulation, where the probability of firing was up-regulated 1 theta cycle (∼110 ms) after a spike (Figure 2a). The first cluster showed suggestive theta modulation at a slower frequency. Measures of burstiness (immediate post-spike interval value) and theta modulation are summarized by cluster in Table 2. The clustering analysis corroborates previous data suggesting that CA1 pyramidal neurons organize into functionally different subclasses (Mizuseki et al. 2011), although it was thought that these subclasses of pyramidal neurons were based on location in the CA1 (superficial vs deep) whereas our clusters can be located within the span of one tetrode. The relative proportions of these two clusters of cells differed by group (Figure 4), with control animals having significantly more refractory cells than either MAM or MAM-E: 6.2% of cells in controls vs 3.5% in both MAM and MAM-E animals. We hypothesize that this relative increase in these less-theta modulated cells in controls may be neurons that have not been recruited into the network for the behavioral task being performed at the time of recording. However, these cells may be available for recruitment if the task becomes harder and therefore could represent a level of redundancy in control animals.

**Table 2.** Description of the PSF parameters of all recorded populations of pyramidal neurons.

|  |  | Immediate Post-Spike Interval Value | std. error | Theta Correlation | std. error | Theta Peak | std. error | Theta Depth | std. error | Theta Trough | std. error |
| --- | --- | --- | --- | --- | --- | --- | --- | --- | --- | --- | --- |
| Bursty | Control | 3.01E-03 | 1.98E-04 | 3.64E-01 | 1.78E-02 | 5.72E-02 | 2.29E-03 | 1.27E-01 | 9.87E-03 | -1.39E-02 | 9.09E-03 |
|  | MAM | 3.13E-03 | 1.63E-04 | 3.64E-01 | 1.95E-03 | 5.21E-02 | 6.12E-04 | 1.26E-01 | 8.30E-04 | -1.40E-02 | 4.44E-04 |
|  | MAM-E | 3.37E-03 | 1.83E-04 | 3.84E-01 | 1.78E-03 | 5.72E-02 | 8.42E-04 | 1.36E-01 | 1.58E-03 | -1.89E-02 | 1.71E-03 |
| Refractory | Control | -3.41E-04 | 3.21E-04 | 3.38E-01 | 4.13E-02 | 3.72E-02 | 5.53E-03 | 1.68E-01 | 6.96E-03 | -4.18E-02 | 5.19E-03 |
|  | MAM | -1.85E-04 | 5.17E-04 | 3.77E-01 | 2.79E-02 | 4.36E-02 | 8.88E-03 | 1.69E-01 | 1.82E-02 | -3.76E-02 | 1.49E-02 |
|  | MAM-E | 2.34E-04 | 5.19E-04 | 3.22E-01 | 4.42E-02 | 7.24E-02 | 3.08E-03 | 1.57E-01 | 1.12E-02 | -1.97E-02 | 7.43E-03 |

**Figure 2.**
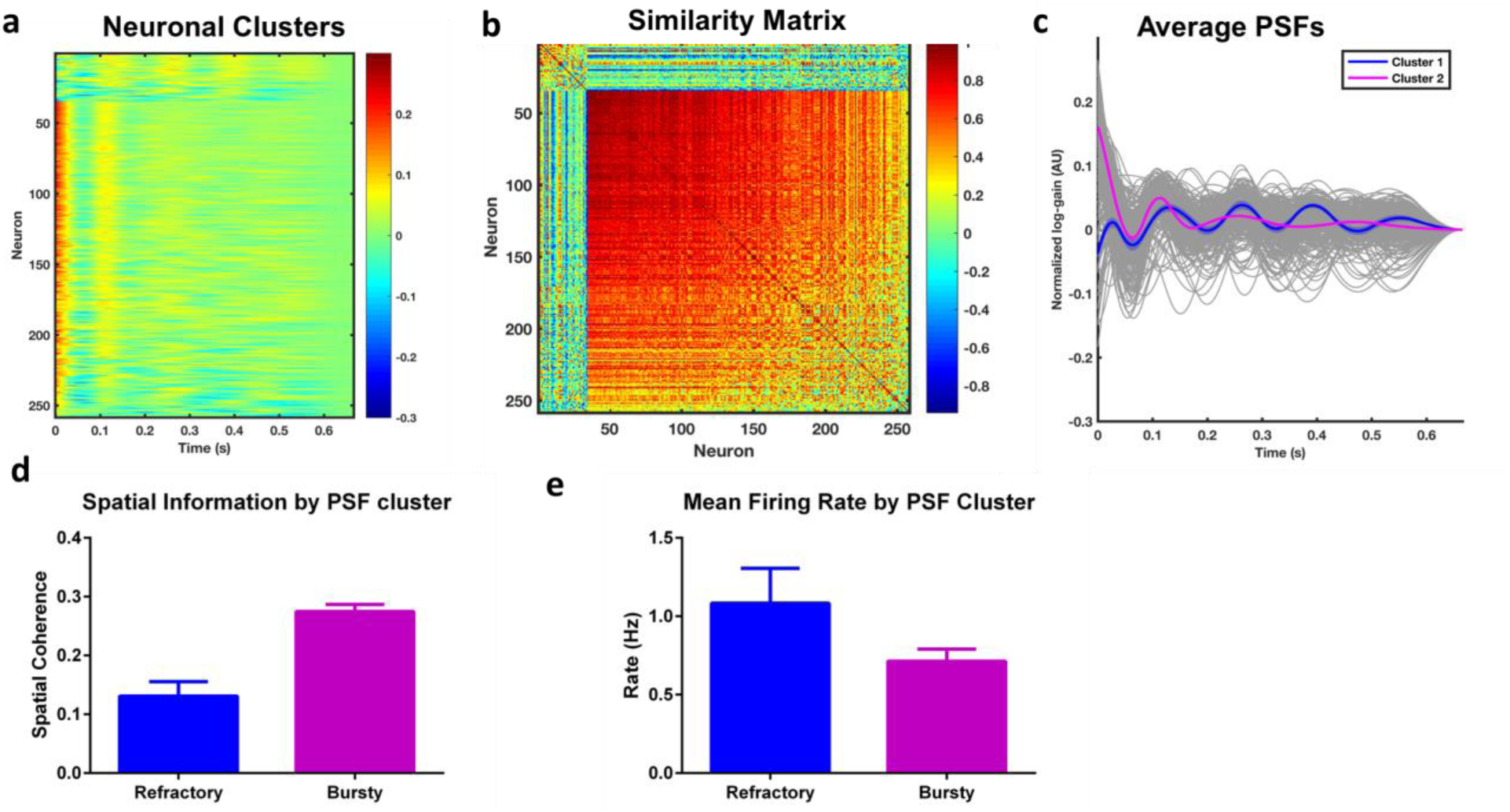
CA1 neurons *in vivo* cluster into two distinct populations, whose distributions differ by group. Neurons from all groups cluster into two main clusters, shown by the heat map of PSFs from the whole population (**a**) sorted by similarity (see also **b**). Average traces of the PSF from bursty and refractory neurons show that the probability of a spike in bursty neurons (**c**; magenta) is rapidly up-regulated in the first 30ms post-spike, followed by down-regulation and subsequent upregulation at 110 ms, approximately one cycle of theta. Refractory neurons do not show this initial upregulation (**c**; blue). Envelopes around the PSFs represent one standard error; gray traces represent individual PSFs for the entire population (**c**). The clusters also contain different amounts of spatial information (**d**), as shown by average spatial coherence of the different cell types, with bursty cells showing significantly higher average coherence. Mean firing rates for the entire session on average are not significantly different between the clusters (**e**).

**Figure 3.**
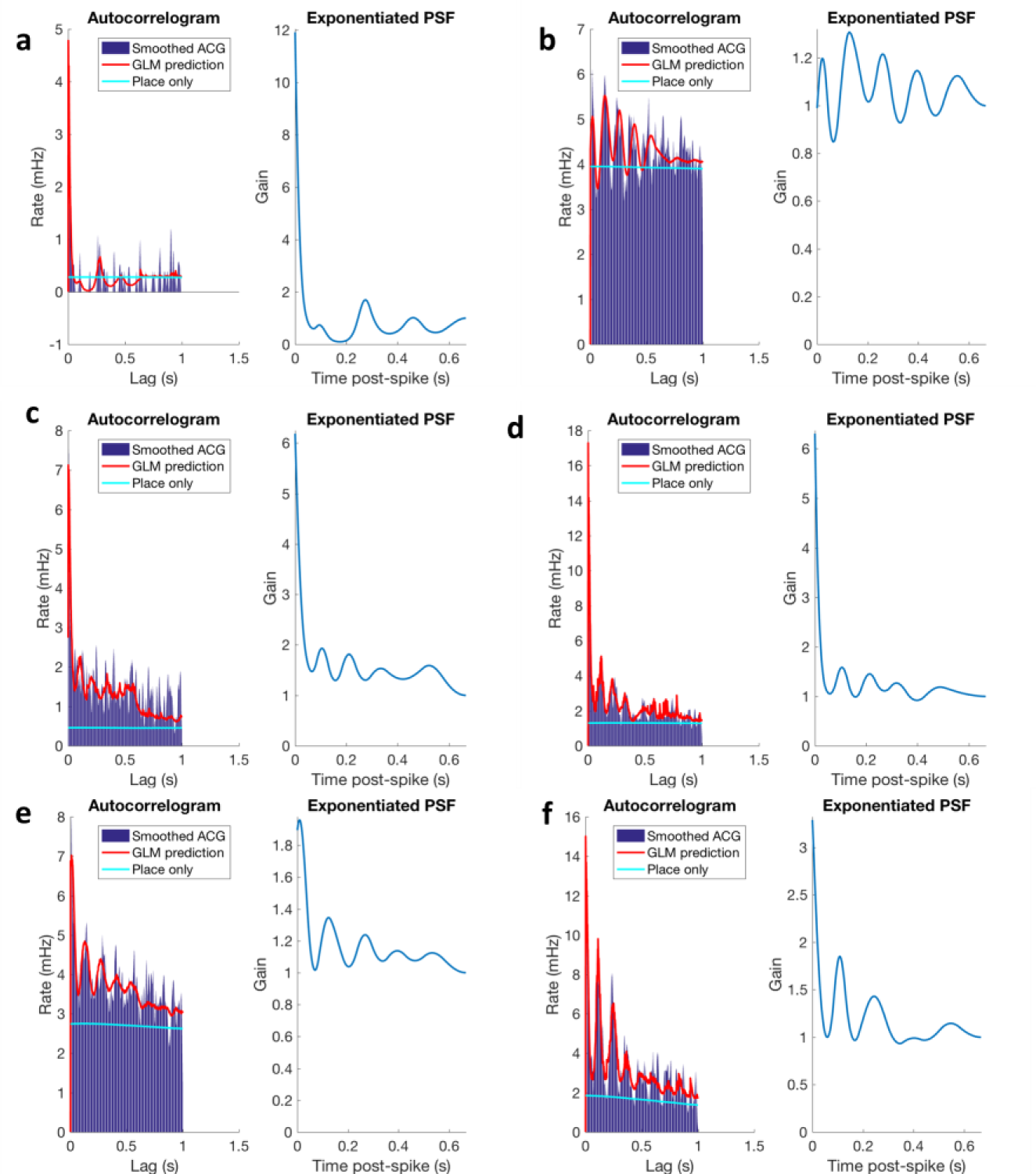
The autocorrelograms (left in each panel) contain oscillatory structure that cannot be modeled with a place field alone (cyan curve). The inclusion of the PSF in the GLM allows the model to accurately predict the autocorrelation structure of neural firing. The exponentiated PSF (right in each panel) corresponds to the time-dependent gain of firing rate. For example, in panel F the neuron has an approximately 3.5-fold up-regulation of firing probability immediately after a spike, and an approximately 1.8-fold up-regulation one theta cycle after a spike. These examples were chosen at the 5^th^, 20^th^, 40^th^, 60^th^, 80^th^, and 95^th^ percentiles of the distribution of coherences (low-to-high; panels a-f respectively).

**Figure 4.**
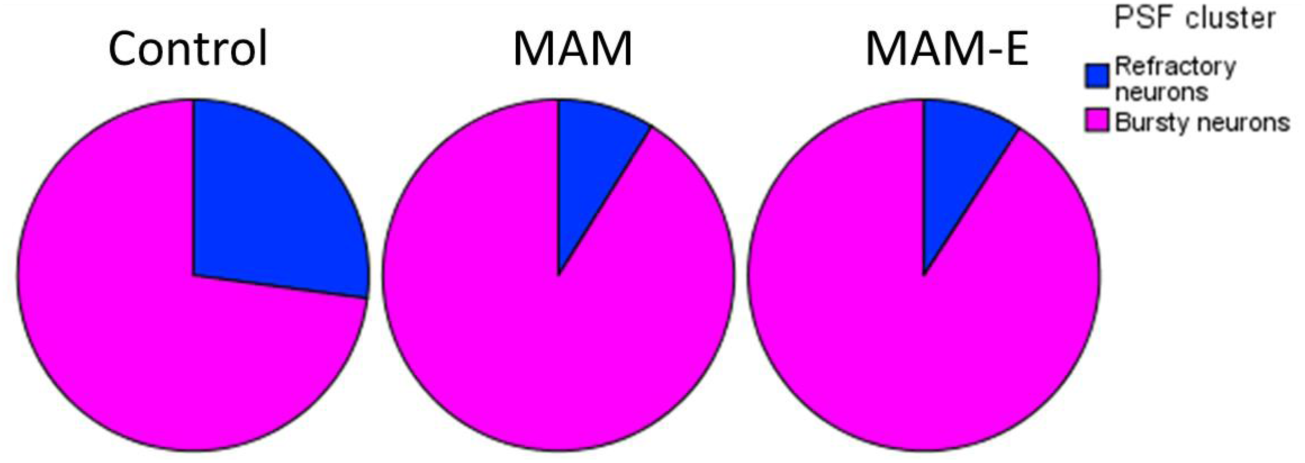
The distribution of the two PSF clusters differs among the groups. Pie charts of bursty (magenta) and refractory (blue) populations shows that the distribution of these neuronal populations differs among the groups, suggesting that post-weaning environmental enrichment does not simply alter the proportion of neurons in the hippocampus with bursty or refractory firing properties.

The spatial coherence of the cells was dramatically different between the clusters (Figure 2d). The bursty cells had significantly higher coherence than the other cells (0.27 ± 0.012 vs. 0.13 ± 0.027, *p* < 0.0001). Because of the critical differences in timing and cognitive content of the bursty cells, we retained only these cells for further analysis.

The PSFs of bursty cells differed between groups (Figure 5a). The MAM and MAM-E cells have significantly higher up-regulation immediately post-spike, while the MAM-E and control cells had stronger up-regulation one theta cycle after a spike. Controls cells showed the strongest theta modulation, with significant up-regulation even at the second theta cycle post-spike.

**Figure 5.**
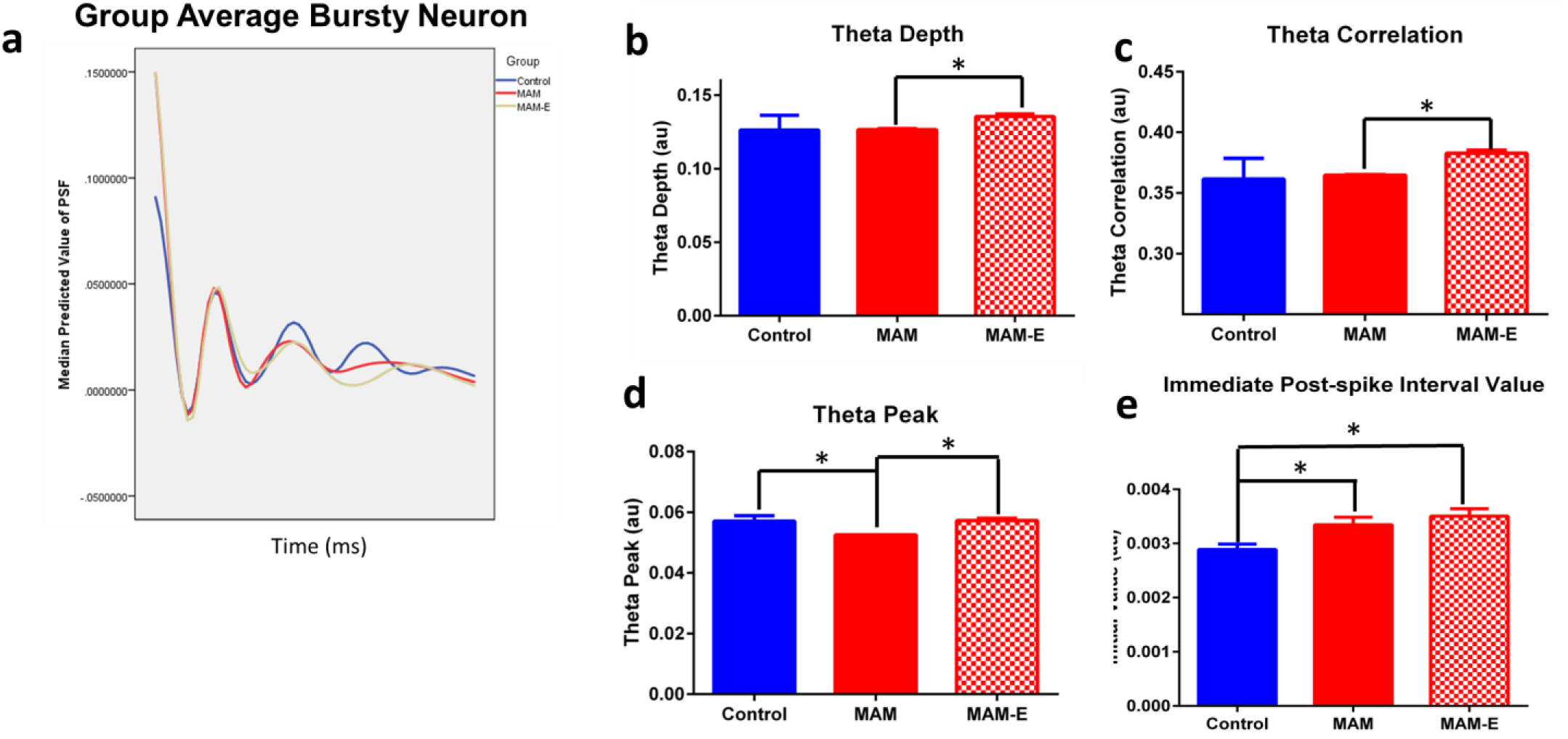
PSF parameters are altered in MAM animals and modified by environmental enrichment. Average post-spike filters by group (**a**) show clear differences in the shape of the filters by group, particularly at the initial ms post-spike, and at timepoints 0.1s post-spike and 0.2s post-spike, corresponding to the first and second phase of theta. MAM neurons (red filled bars) reduced theta depth (**b**) and reduced theta correlation (**c**) compared to MAM-E animals (red checkered bars). MAM neurons also show reduced theta peak (**d**) compared to both MAM-E and control animals (blue bars). MAM and MAM-E neurons both show increased burstiness (**e**) compared to controls.

### Animals with MCD exhibit altered timing coding that is partially restored by EE

To quantify differences in PSFs among groups and make associations between FST and spatial coherence, we parameterized the PSFs by key features that captured the significant quantitative variation in population. The first parameters we examined were modulation of firing in the theta frequency range (6-12 Hz). In well-modulated neurons the theta trough should be low (down-regulation out of phase with theta) and the peak probability of firing should be high (up-regulation in phase with theta). Neurons from MAM animals have less precise theta modulation (Figure 5b-d; p<0.001) with a higher theta trough, lower theta correlation and lower theta depth when compared to MAM enriched animals (see Table 2 for details).

In addition to modulation within the theta frequency range, we observed differences in the up-regulation of firing in MCD animals immediately after a spike. We defined the *burstiness* of a PSF as the integral of the normalized PSF over the interval 0-30 ms (Figure 2c). This captures the strength of up-regulation immediately after a spike, i.e. increased burstiness. Consistent with previous in-vitro experiments showing increased bursting and hyperexcitability in MAM animals (Baraban & Schwartzkroin 1995) both MAM and MAM-E animals have increased initial value compared to controls (0.0029 ± 0.0001 in controls vs 0.0033 ± 0.0001 in MAM and 0.004 ± 0.0001 in MAM-E, *p* < 0.01; Figure 5e). The significant difference between MAM-E and control cells in this initial interval suggests that some timing coding changes persist in brains with MCDs, even after rescue of behavioral and cognitive phenotypes.

### Variation in fine spike timing predicts spatial coherence

The PSFs measure timing properties of neuronal firing while spatial coherence measures the quality of the neural encoding of external space. The former derive from local circuit dynamics, while the latter measures the match between the outside world and its internal representation. Significantly, however, the PSF parameters predict spatial coherence. Group, initial value, theta depth, theta correlation, and theta peak height were correlated with coherence in a multivariate regression that included the main effects and all two-way interactions. Remarkably, the PSF parameters accounted for 58.5% of the variance in spatial coherence in this regression (Pearson’s r^2^ = 0.585, p<0.001; Figure 7c), suggesting that the fine spike timing of the CA1 pyramidal cell firing is directly related to place-modulated rate coding of these neurons.

To corroborate the above findings, and determine whether we identified a comprehensive set of timing parameters, we performed principal component analysis (PCA) on the set of PSFs. Because we used a 10-dimensional basis for the PSFs (see Materials and Methods) there were ten factors that explained 100% of the variance in the PSFs (Figure 6). Using these 10 factors as input to a multivariate regression (including pairwise interaction terms), we found that three factors were significantly associated with coherence (true vs. predicted R^2^ = 0.615; Figure 7d). These factors had strong loadings in the interval just after the spike, and at the first and second theta cycles post-spike (Figure 6), demonstrating that the critical time points for predicting coherence align with the immediate post-spike interval and the peak of each subsequent theta cycle. Moreover, the scores for each of these factors differed among the experimental groups. The 10 factors were compared across groups and validate the findings from analyses of the *a priori* defined parameters of the PSF (Figure 6). Importantly, the significant factors that corresponded to theta modulation were higher in controls and MAM-E, whist the significant factor that corresponded to out of theta firing was higher in MAM animals, further supporting the view that MAM animals have disrupted timing coding (Figure 6). This indicates that these time points are differentially regulated among the groups.

**Figure 6.**
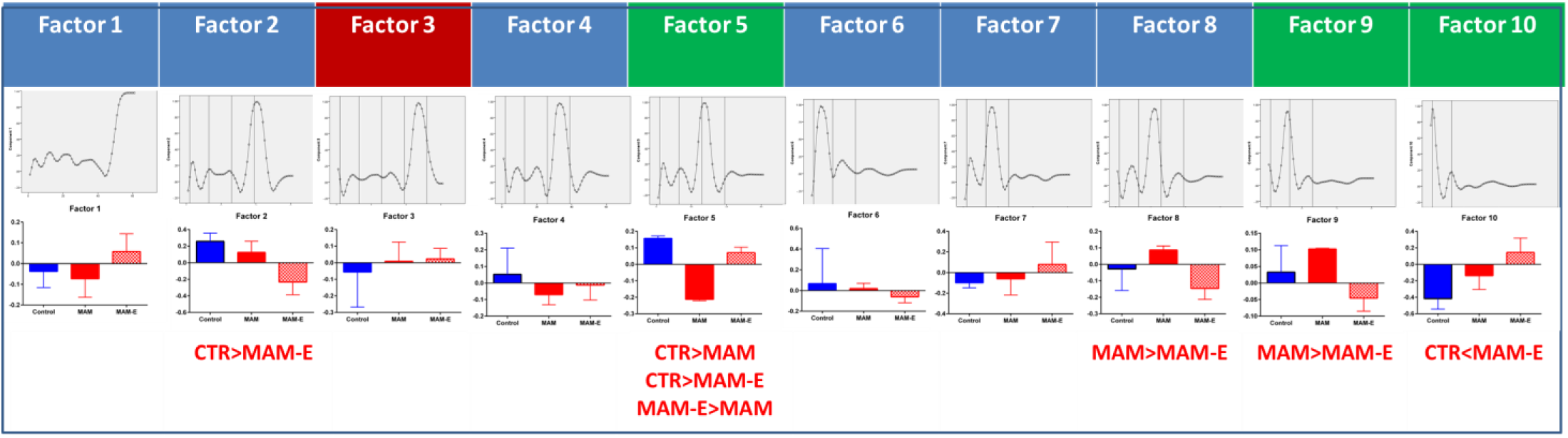
PSF factors are altered in MAM animals and modified by environmental enrichment. A PCA divides the PSF into 10 factors, shown as loading plots for each of the individual factors in the top row. Vertical lines represent the initial value time point (first line) and successive oscillatory cycles of theta (second-fourth lines). Factors 3, 5, 9 and 10 significantly predict spatial coherence; factors 5, 9 and 10 positively predict coherence (green label) while factor 3 negatively predicts coherence (red label; p>0.001). Factors 2, 5, 8, 9 and 10 show group differences. Differences between MAM-E and control cells in factor 2 suggest that MAM-E cells are less modulated at the third theta peak; however, this factor is not significantly related to spatial coherence. Differences between MAM and MAM-E cells in factor 8 suggest that there is more modulation in between theta peaks in MAM cells; however, this factor is also not significantly related to spatial coherence. Factors 5, 9 and 10 relate to spatial coherence and show group differences as well. These factors show strong modulation at the second (factor 5) and first (factor 9) theta peaks, as well as at the initial post-spike interval region (factor 10), thus corroborating the importance of fine spike timing in these time periods.

**Figure 7.**
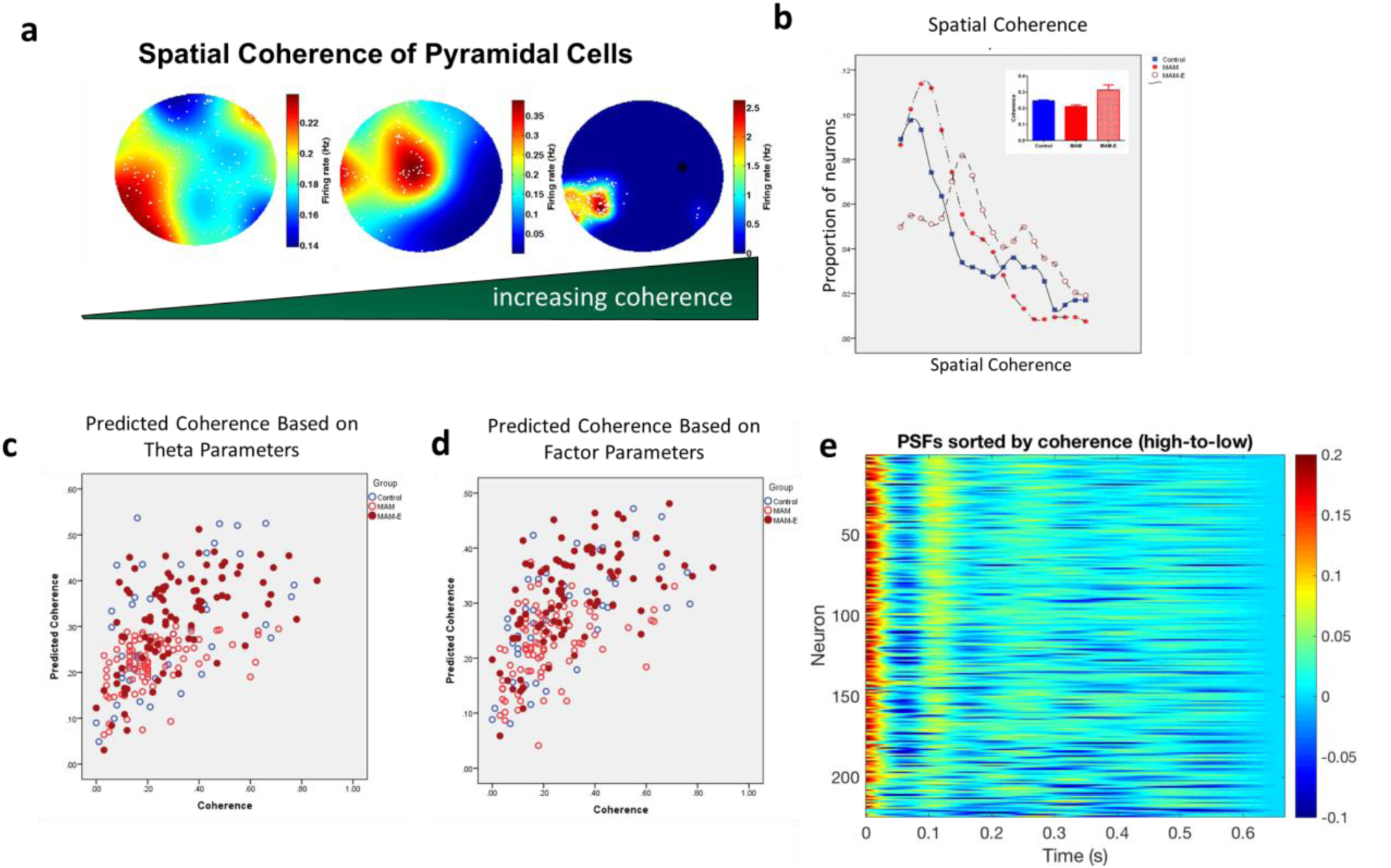
PSF parameters predict spatial coherence. Spatial coherence is a good measure of how well a location in space is rate encoded (**a**); cells with higher coherence (right) have very tight regulation of firing in a specific region of space, whereas cells with lower coherence (left) show more promiscuous firing with very little rate modulation throughout the arena and many out of field action potentials (shown as white dots). Panel (**b**), shows the spatial coherence values as a moving average of four from values of 0 to 0.9 between the groups. This plot illustrates that the coherence distributions are shifted to the right (towards higher coherence values) in the MAM-E group (open red circles) compared to the controls (closed blue circles) or the MAM animals (closed red circles). Control coherence values have a longer and higher right tail, this tail appears to be absent in the MAM group. This is also seen in plots of group average coherence of pyramidal cells (**b**, inset) in MAM animals (red bars) compared to controls (blue bars); this is restored with enrichment (red checkered bars). These measures of fine-spike timing do not, necessarily, have to correlate to spatially-modulated rate coding of pyramidal neurons in the hippocampus; however predicted coherence, obtained from regression analyses of the PSF parameters against coherence, plotted against actual coherence shows a clear positive relationship (**c;** R^2^ = 0.565) between PSF parameters and spatial coherence in all groups. This confirms that the PSF parameters predict spatial coherence. Likewise, factor scores also significantly positively predict spatial coherence as well (**d;** R^2^= 0.615). Heat map of the PSFs from bursty CA1 pyramidal neurons (d) sorted by coherence high to low, shows that cells at the top of the heat map (high coherence) have greater burstiness shown as warmer colors at the initial time points and cells as the bottom of have greater total variation in the post-spike filter, in particular more shallow valleys before the first peak of theta at 0.1-0.2s and more modulation after one theta cycle after 0.2s.

### EE is associated with a partial rescue of parvalbumin-positive interneurons in CA1

Interneurons, particularly somatic and axon initial segment-targeting interneurons, are crucial for timing of pyramidal cell outputs(Klausberger et al. 2003) This subset of interneurons often expresses the calcium-binding protein marker parvalbumin (PV). Given the aberrant temporal coding in MAM animals and restoration MAM-E animals, we examined counts of PV^+^ neurons post-mortem. We found a stark reduction in PV^+^ neurons in MAM adult animals from 4154.89 ± 911.05 cells/10,000 μm^2^ in controls to 1747.23 ± 62.58 cells/10,000 μm^2^ (p<0.0001, Figure 8). Enrichment attenuated this decrease, with MAM-E animals 2363.35 ± 219.49 cells/10,000 μm^2^. Given the importance of PV interneurons for temporal organization of pyramidal neuron firing, we suggest that this may represent a major cellular mechanism by which enrichment improves temporal structure.

**Figure 8.**
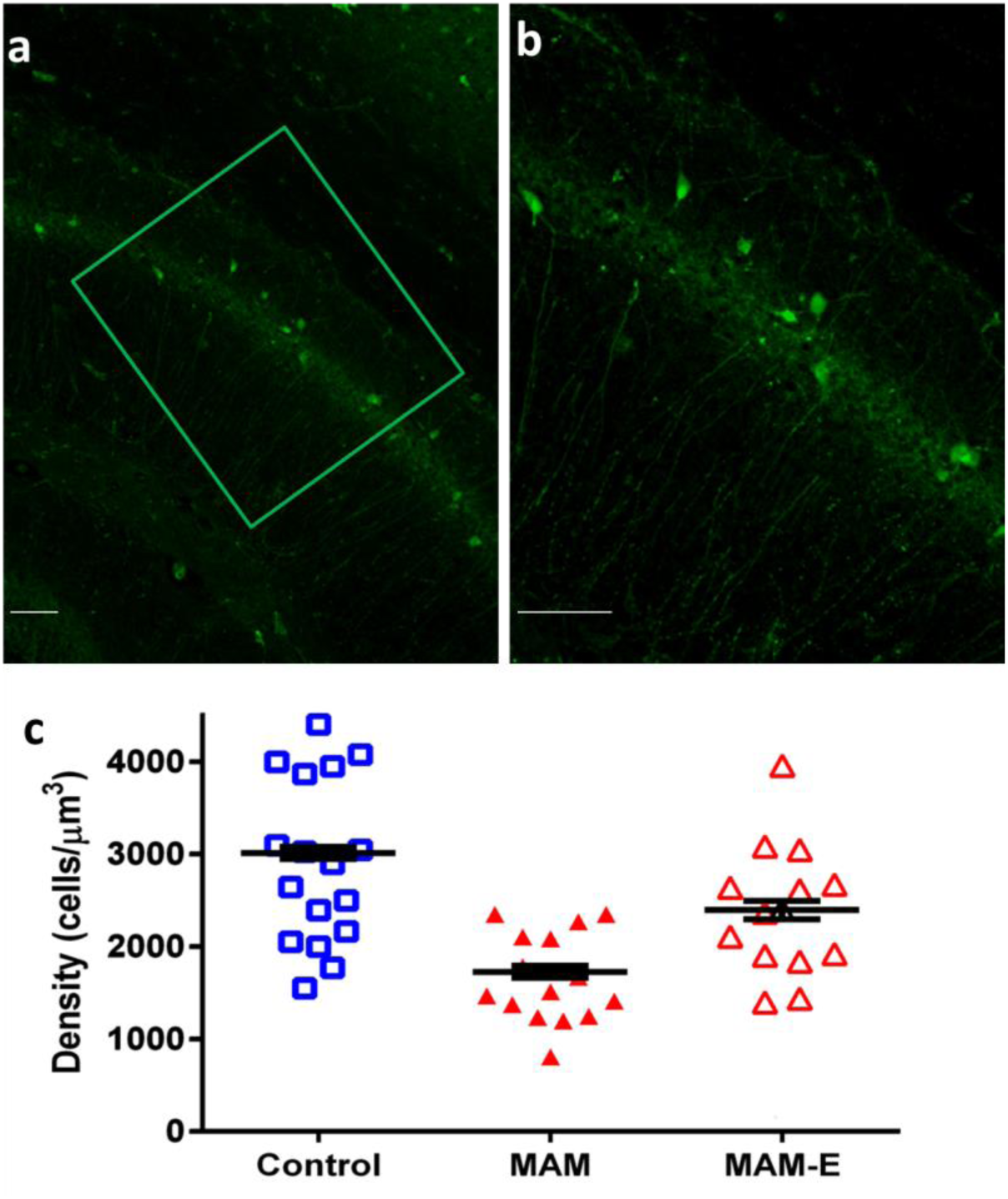
Environmental enrichment normalizes PV^+^ interneuron numbers. Immunohistochemistry for parvalbumin positive interneurons at 10× (**a**) and 20× (**b**) magnification. Scale bars are 100 μm. Quantification (**c**) in MAM animals (red closed triangles) show a stark decrease in the number of PV^+^ interneurons per μm^3^ in CA1 compared to their control counterparts (blue squares). Environmental enrichment (red open triangles) is able to significantly increase the numbers of PV^+^ interneurons in CA1.

## DISCUSSION

Our results definitively show that rats with MAM-induced malformations of cortical development have disruptions to hippocampal rate and timing coding that likely underpin previously identified impairments in spatial cognition. Despite the presence of a significant structural brain abnormality, exposure of MAM rats to post-weaning environmental enrichment restores several characteristics of the neural dynamics towards normal, consistent with behavioral observations. Importantly, there is a clear relationship between the timing fidelity of action potential firing, as measured with a post-spike filter, and the spatial fidelity as measured by spatial coherence. Environmental enrichment is the only therapeutic technique that has been shown to improve cognition in any model of MCDs, so it presents a critical window into the functional changes that must occur in response to successful treatment. In this way, our data suggest that disrupted neural dynamics can be conceptualized as a system-level mechanism of cognitive impairment. Therefore, therapies that directly target neural dynamics have enormous promise for ultimately improving cognitive outcomes in patients with malformations of cortical development.

The hippocampus is a crucial brain structure supporting cognition and is a site of massive integration of information. Healthy brain function requires neural activity to be precisely timed, allowing effective coordination within and between brain regions (Buzsáki 2002). When the hippocampus is damaged or malformed there are a huge number of alterations to brain physiology, including changes in neuron number (Penschuck et al. 2006), synaptic function (Calcagnotto 2005), a variety of cell signaling pathways including neurotrophic pathways (Fiore et al. 2004) and neurotransmission (Snyder et al. 2013; Hradetzky et al. 2012), all of which could potentially contribute to impaired cognition, independently or in concert. Restoring cognition post-insult requires modifying the pathological dynamics of the impaired neural network. In the case of MCDs, the brain must generate sufficiently normal-like dynamics using an abnormal neural substrate. Identifying therapeutic targets to induce such changes in dynamics is critical for developing therapies for improving cognitive outcomes for patients with MCDs. These targets need to capture the function of the full neural network as a complex system, and necessarily exist at an abstract layer above all of the genetic, molecular, and cellular processes that are critical to maintain a living neural network. Our results are a significant step in this direction. We show that the encoding of external spatial information, the cognitive representation, is strongly related the timing properties of firing, and that this quantitative relationship persists across control animals, animals with MCDs, and animals that received EE. In other words, timing coding predicts cognitive encoding quality across normal and abnormal brains, and in the presence or absence of cognitive impairment. We posit, therefore, that timing coding itself is a systems-level target for improving cognitive outcome in MCDs.

The MAM model in our hands is used to model an etiology associated with intractable pediatric epilepsy. However, *in utero* insults and cortical malformations are common to many neurodevelopmental disorders (Cascella et al. 2009; Piontkewitz et al. 2011; Casanova et al. 2013). Therefore, understanding how abnormal neural networks can reorganize to support optimal cognition may have far-reaching implications. In prior work, we have shown that EE in MCDs leads to cognitive gains (Jenks et al. 2013). In the present work, we take an important step further to show that EE restores multiple features of timing coding and that these alterations predict improved cognitive encodings. Moreover, this improvement with EE is concurrent with the preservation of at least one class of neurons (PV^+^ interneurons) that primarily control the spike timing of CA1 pyramidal cells through dynamically modulated somatic inhibition. While we do not believe that increases in PV^+^ interneuron numbers are the only relevant histological change in MAM-E brains, there is a well-established relationship between these neurons and the theta oscillatory behavior of CA1 pyramidal cells(Klausberger et al. 2003), making them a highly likely component of the substrate producing the systems-level changes we observe in spike timing.

The direct relationship between timing and cognition that we found spanning both a disease model (MAM) and a therapeutic rescue model (MAM-E) suggests that modifying neural dynamics is critical for restoring cognition. Our results are thus complementary to important recent work showing that driving neural circuits at specific frequencies can result in cognitive gains in patients with Alzheimer’s disease and other diseases (Kuo et al. 2014). Our results suggest the development of therapies that directly intervene in the timing coding of cells. The advent of sophisticated electrical and optogenetic stimulation approaches, along with cell-based therapies such as interneuron precursor implantation, makes such strategies potentially viable.

## AUTHOR CONTRIBUTIONS

AEH, WC, AM, MML, GR collected data. AEH, JMM and RCS analyzed data. AEH, JMM, GLH, RCS wrote paper.

